# Mannose receptor is a restriction factor of HIV in macrophages and is counteracted by the accessory protein Vpr

**DOI:** 10.1101/742197

**Authors:** Jay Lubow, David R. Collins, Mike Mashiba, Brian Peterson, Maria Virgilio, Kathleen L Collins

## Abstract

HIV-1 Vpr is necessary to support HIV infection and spread in macrophages. Evolutionary conservation of Vpr suggests an important yet poorly understood role for macrophages in HIV pathogenesis. Vpr counteracts a previously unknown macrophage-specific restriction factor that targets and reduces the expression of HIV Env. Here, we report that the macrophage mannose receptor (MR), is the restriction factor targeting Env in primary human monocyte-derived macrophages. Vpr acts synergistically with HIV Nef to target distinct stages of the MR biosynthetic pathway and dramatically reduce MR expression. Silencing MR or deleting mannose residues on Env rescues Env expression in HIV-1-infected macrophages lacking Vpr. However, we also show that disrupting interactions between Env and MR reduces initial infection of macrophages by cell-free virus. Together these results reveal a Vpr-Nef-Env axis that hijacks a macrophage mannose-MR response system to facilitate infection while evading MR’s normal role, which is to trap and destroy mannose-expressing pathogens.

## Introduction

Vpr is a highly conserved HIV accessory protein that is necessary for optimal replication in macrophages (Balliet, Kolson et al. 1994) but its mechanism of action is poorly understood. Studies using human lymphoid tissue (HLT), which are rich in both T cells and macrophages, have found that loss of Vpr decreases virus production (Rucker, Grivel et al. 2004) but only when the virus strain used is capable of efficiently infecting macrophages (Eckstein, Sherman et al. 2001). These studies provide evidence that Vpr enhances infection of macrophages and increases viral burden in macrophage containing tissues. Because Vpr is packaged into the virion (Cohen, Dehni et al. 1990) and localizes to the nucleus (Lu, Spearman et al. 1993), it may enhance early viral replication events. However, mononuclear phagocytes infected with viral particles in which Vpr is provided by trans-complementation in the producer cells still have a *vpr*- null phenotype (Connor, Chen et al. 1995), indicating that Vpr’s role in the HIV replication cycle continues into late stages.

Previous work by our group demonstrated that Vpr counteracts an unidentified, macrophage-specific restriction factor that targets Env and Env-containing virions for lysosomal degradation (Mashiba, Collins et al. 2014, Collins, Lubow et al. 2015). This restriction could be conferred to permissive HEK293T cells by fusing them with MDM to create HEK293T-MDM heterokaryons. A follow up study demonstrated that by increasing steady state levels of Env, Vpr increases formation of virological synapses between infected MDM and autologous, uninfected T cells (Collins, Lubow et al. 2015). This enhances spread to T cells and dramatically increases levels of Gag p24 in the culture supernatant. This finding helps explain the paradoxical observations that Vpr is required for maximal infection of T cells in vivo (Hoch, Lang et al. 1995) but numerous studies have shown Vpr only marginally impacts infection of pure T cell cultures in vitro [e.g. (Mashiba, Collins et al. 2014)].

Our goal in the current study was to identify and characterize the myeloid restriction factor targeting Env that is counteracted by Vpr. We reasoned that macrophage-specific Env-binding proteins, including the carbohydrate binding protein mannose receptor (MR), were candidates. MR is expressed on several types of macrophages *in vivo* (Liang, Shi et al. 1996, Linehan, Martinez-Pomares et al. 1999) and is known to mediate innate immunity against various pathogens (Macedo-Ramos, Batista et al. 2014, Subramanian, Neill et al. 2019). MR recognizes mannose rich structures including high-mannose glycans, which are incorporated in many proteins during synthesis. In eukaryotic cells most high-mannose glycans are cleaved by *α*-mannosidases and replaced with complex-type glycans as they transit through the secretory pathway. By contrast, in prokaryotic cells, high-mannose residues remain intact, making them a useful target of pattern recognition receptors including MR. Some viral proteins, including HIV-1 Env, evade mannose trimming (Coss, Vasiljevic et al. 2016) and retain enough high-mannose to bind MR (Trujillo, Rogers et al. 2007, Lai, Bernhard et al. 2009). There is evidence that HIV-1 proteins Nef and Tat decrease expression of MR based on studies performed in monocyte derived macrophages (MDM) and the U937 cell line, respectively (Caldwell, Egan et al. 2000, Vigerust, Egan et al. 2005). Nef dysregulates MR trafficking using an SDXXLΦ motif in MR’s cytoplasmic tail (Vigerust, Egan et al. 2005), which is similar to the sequence in CD4’s tail that Nef uses to remove it from the cell surface (Bresnahan, Yonemoto et al. 1998, Greenberg, DeTulleo et al. 1998, Cluet, Bertsch et al. 2005). Whether MR or its modulation by viral proteins alters the course of viral replication has not been established.

Here, we confirm that Nef reduces MR expression in primary human MDM, although in our system, the effect of Nef alone was relatively small. In contrast, we report that co-expression of Vpr and Nef dramatically reduced MR expression. In the absence of both Vpr and Nef, MR levels normalized indicating that Tat did not play a significant, independent role in MR downmodulation. Deleting mannose residues on Env or silencing MR alleviated mannose-dependent interactions between MR and Env and reduced the requirement for Vpr. Although the post-infection interactions between MR and Env reduced Env levels and inhibited viral release, we provide evidence that these same interactions were beneficial for initial infection of MDM. Together these results reveal mannose residues on Env and the accessory proteins Nef and Vpr are needed for HIV to utilize and then disable an important component of the myeloid innate response against pathogens intended to thwart infection.

## Results

### Identification of restriction factor counteracted by Vpr in primary human monocyte-derived macrophages

Because we had previously determined that Vpr functions in macrophages to counteract a macrophage selective restriction factor that targets Env, we reasoned that Env-binding proteins selectively expressed in macrophages were potential candidate restriction factors. To determine wether any factors fitting this description were targeted by Vpr, we performed western blot analysis of candidate Env binding proteins in MDM matched for wild type or Vpr null HIV infection frequency (Figures 1A and B). We found that one such protein, mannose receptor (MR), which is highly expressed on macrophages and has been previously shown to bind Env (Trujillo, Rogers et al. 2007, Fanibunda, Velhal et al. 2008, Lai, Bernhard et al. 2009), was significantly decreased by wild type HIV 89.6 but not by 89.6 *vpr*-null (Figure 1C and D, *p<*0.01).

**Figure 1:**
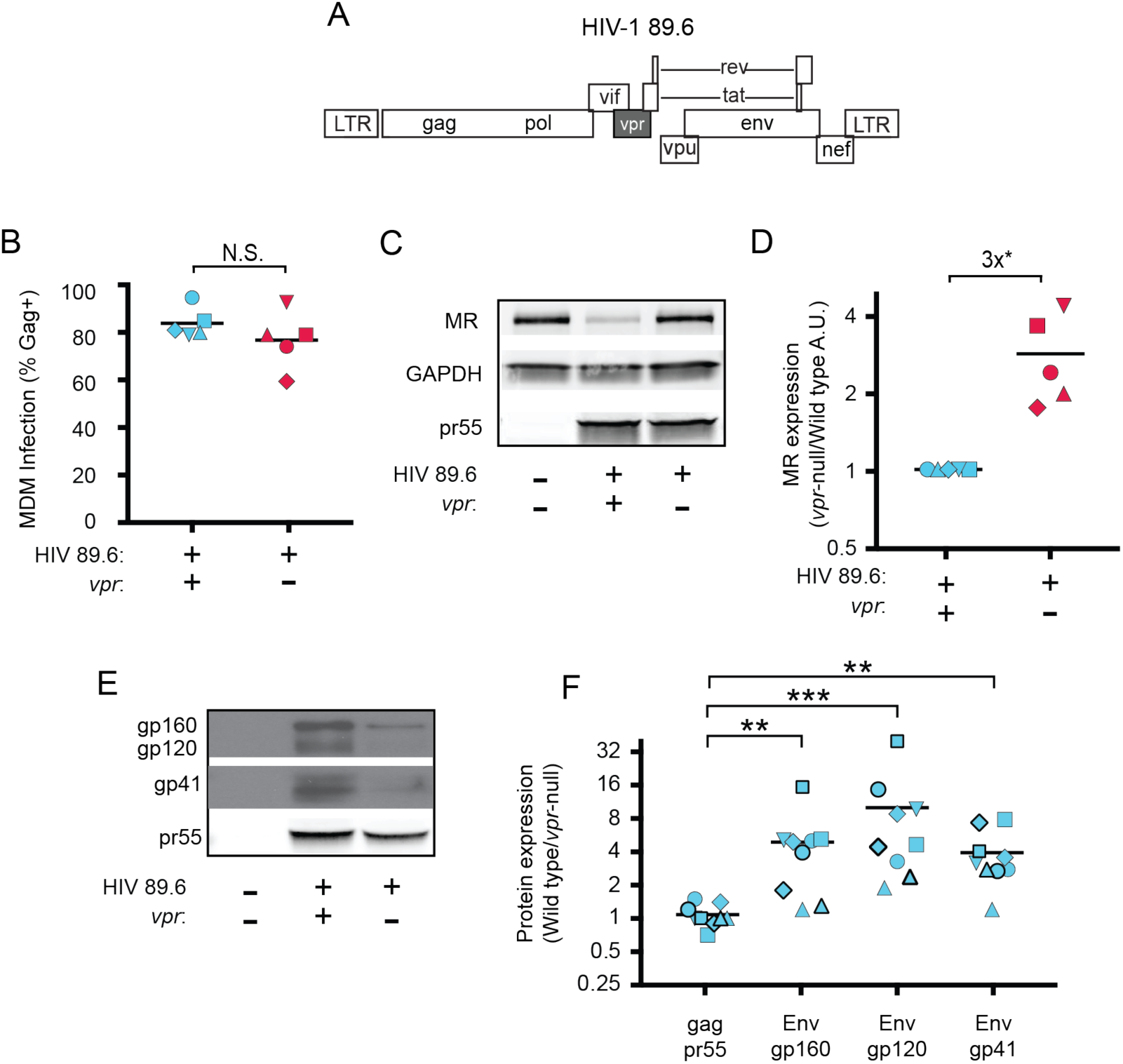
HIV Vpr reduces steady state levels of host mannose receptor in MDM and increases steady state levels of viral Env protein. A) Diagram of the HIV 89.6 proviral genome. The shaded box shows the location of *vpr,* which was disrupted by a frame shift mutation to create the Vpr-null version (Mashiba, Collins et al. 2014). HIV-1 89.6 is a dual CXCR4/CCR5-tropic HIV molecular clone isolated from the peripheral blood of an AIDS patient (Collman, Balliet et al. 1992). B) Summary graph depicting MDM infected by HIV 89.6 wild type and *vpr*-null with matched infection frequencies of at least 50% 10 days post infection as measured flow cytometrically by intracellular Gag p24 staining. This subset with high frequencies of infection was selected to examine potential effects on host factors. Statistical significance was determined using a two-tailed, paired *t*-test. N.S. – not significant, p=0.34 C) Western blot analysis of whole cell lysates from MDM prepared as in B. D) Summary graph displaying relative expression of MR in wild type and mutant 89.6 from blots as shown in B. Western blot protein bands were quantified using a Typhoon scanner. Values for MR expression in MDM infected with Vpr-null HIV were normalized to GAPDH and then to wild type for each donor. Statistical significance was determined using a two-tailed, paired *t*-test. * p=0.03 E) Western blot analysis of HIV protein expression in MDM infected as in B. F) Summary graph of HIV protein expression from western blot analysis as in E and quantified as described in methods. The ratio of expression in wild type to *vpr*-null infection is shown. Data from 9 independent donors with similar frequencies of infection (within 2-fold) following ten days of infection are shown. Statistical significance was determined using a two-tailed, ratio *t*-test, ** p<0.01, *** p<0.001. Data from each donor is represented by the same symbol in all charts. Mean values are indicated.

To confirm the effect of Vpr on Env during infections or primary human macrophages in which MR was downmodulated, we performed quantitiative western blot analysis. As shown in Figures 1E and F, we confirmed that amounts of Vpr sufficient for MR downmodulation were also sufficient for stabilizing expression of Env (gp160, gp120, gp41). Compiled data from nine donors clearly demonstrated results that were similar to our prior publication (Mashiba, Collins et al. 2014); under conditions of matched infection in which there was no significant difference in pr55 levels between wild type and *vpr*-null infections, all three forms of Env were significantly more abundant in the wild type infection (gp160: 4-fold, *p*<0.002; gp120: 6-fold, *p*<0.002; gp41: 4-fold, *p* <0.001).

### Combined effects of Nef and Vpr have dramatic effects on MR levels in a subset of infected cells

Because an earlier report indicated that Nef decreases surface expression of MR (Vigerust, Egan et al. 2005), we asked whether Nef was playing a role in MR downmodulation in our systems. Because HIVs lacking Vpr and Nef spread too inefficiently in MDM to observe effects on host proteins by western blot analysis, we utilized a replication defective HIV with a GFP marker (NL4-3 ΔGPE-GFP) to allow measurement of MR expression via flow cytometry following single-round transduction. This construct has the additional advantage that it eliminates potentially confounding effects of differences in wild type and Vpr-null viral spread. Therefore, we generated the necessary mutations in *nef* and *vpr* and confirmed that these mutations only affected expression of the altered gene product in transfected HEK293T (Figure 2B). For experiments in primary human macrophages, MDM were harvested at earlier times than the experiments described in Figure 1 (five days versus ten days) because of the non-spreading nature of the virus and the capacity to use flow cytometry to identify the subset of infected cells by GFP expression (Figure 2C). Under these conditions, we found that MR expression was dramatically reduced in a subset of GFP^+^ cells when both Vpr and Nef were expressed (Figure 2C-E). Loss of function mutations in either Vpr or Nef led to modest but statistically significant MR downmodulation in a subset of cells. Combined loss of both Nef and Vpr virtually eliminated MR downmodulation (Figure 2D and E). These differences in MR downmodulation were not due to variations in multiplicity of infection of the different viral constructs as MDM transduced with the mutant viral constructs had roughly similar or slightly higher transduction rates as the parental construct (Figure 2F) but demonstrated less MR downmodulation (Figure 2E).

**Figure 2:**
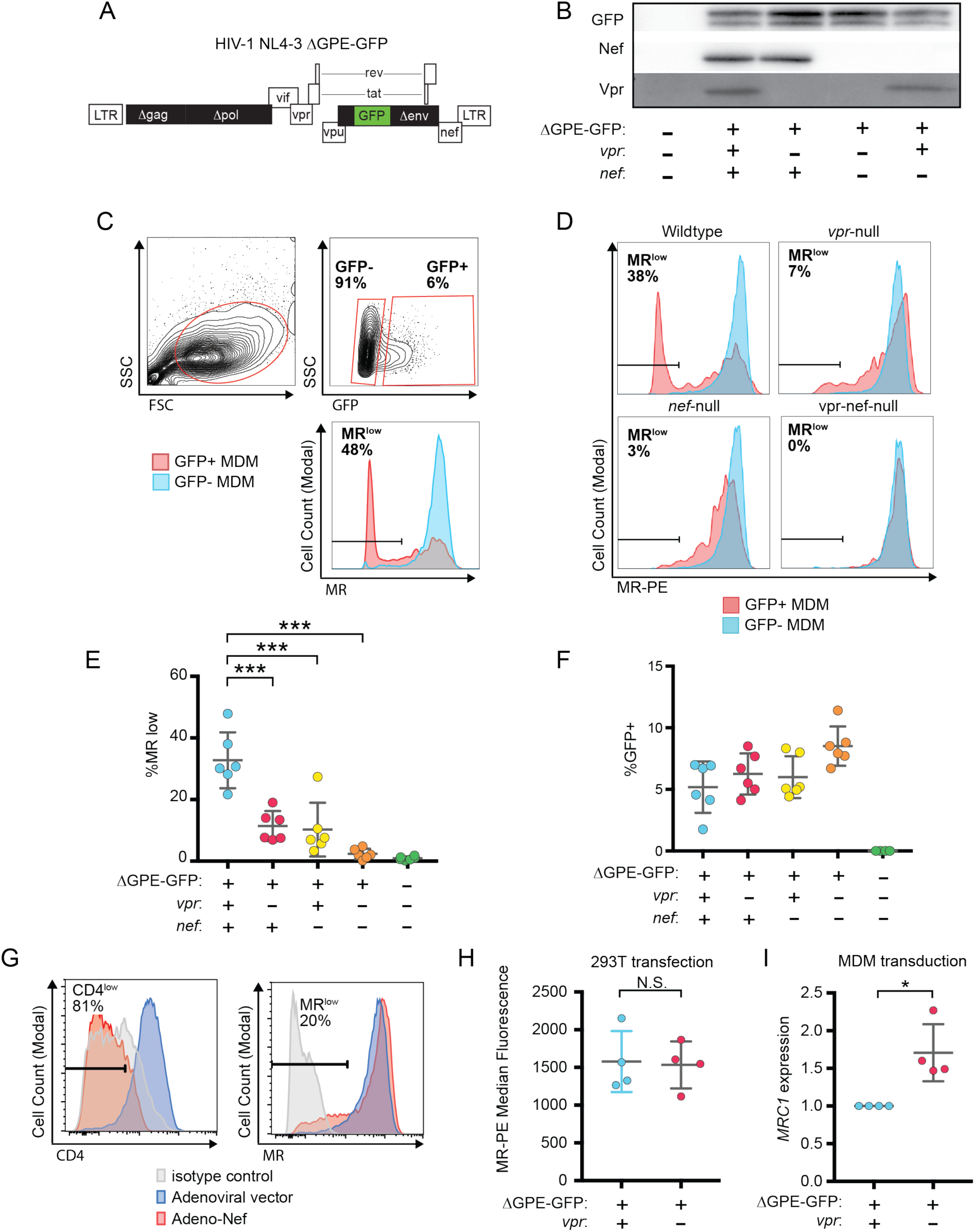
Combined effects of Nef and Vpr completely remove MR from a significant proportion of infected cells at early time points. A) Diagram of HIV NL4-3 ΔGPE-GFP. B) Western blot analysis of whole cell lysates from HEK293T cells transfected with the indicated viral expression construct. C) The gating strategy used to identify live GFP^+^ vs GFP^-^ cells and the fraction of cells that are MR^low^. D) Representative flow cytometric analysis of MDM at five days post transduction by the indicated virus. The percentage of GFP^+^ cells that fell into the MR^low^ gate is indicated in each panel. E) Summary graph depicting the percentage of GFP^+^ cells that fell into the MR^low^ gate in transduced MDM. For the uninfected column the GFP^-^ cells are displayed. (*n=*6 independent donors) F) Summary graph depicting the frequency of transduced (GFP^+^) MDM at the time of harvest. G) Representative flow cytometric plots of MDM transduced with the indicated adenoviral vector (*n*=3 independent donors). H) Summary graph of MR levels as measured by flow cytometry in HEK293T cells co-transfected with the indicated HIV construct. Transfected (GFP^+^) cells were identified using the gating strategy shown in Figure S1. (*n=* four independent transfections) I) Summary graph of mannose receptor (*MRC1*) mRNA expression in MDM transduced with the indicated HIV reporter and sorted for GFP expression by FACS. Expression of *MRC1* was measured by qRT-PCR and normalized to *ACTB* and to the wild type transduction. (*n=*4 independent donors). Mean +/-standard deviation is shown. Statistical significance was determined by a two-tailed, paired *t*-test. N.S. not significant: p=0.89, * p=0.03, *** p<0.001,

To determine whether the modest effect of Nef alone was due to using HIV to deliver Nef as compared to an adenoviral vector delivery system used in a prior publication (Vigerust, Egan et al. 2005), we repeated the experiment using an adenoviral vector expressing Nef. These experiments confirmed that levels of Nef sufficient to downmodulate the HIV receptor, CD4 on nearly all MDM in the culture achieved only modest effects on MR in a subset of cells (Figure 2G) similar to what was observed using the HIV reporter construct (Figure 2E). Thus, Nef and Vpr have modest but significant effects on MR when expressed individually, however the combined effects of both proteins can achieve nearly complete downmodulation at least in a subset of infected cells.

While the effect of Nef has been previously reported and found to be due to disruption of MR intracellular trafficking (Vigerust, Egan et al. 2005), the effect of Vpr on MR has not been previously reported. Vpr is known to target cellular proteins involved in DNA repair pathways for proteasomal degradation via interactions with Vpr binding protein [DCAF1, (McCall, Miliani de Marval et al. 2008)], a component of the cellular DCAF1-DDB1-CUL4 E3 ubiquitin ligase complex (Belzile, Duisit et al. 2007, Hrecka, Gierszewska et al. 2007, Le Rouzic, Belaidouni et al. 2007, Wen, Duus et al. 2007, Lahouassa, Blondot et al. 2016, Wu, Zhou et al. 2016, Zhou, DeLucia et al. 2016). Using this mechanism, Vpr degrades the uracil deglycosylases UNG2 and SMUG1 in HEK293T cells following co-transfection (Schrofelbauer, Yu et al. 2005, Schrofelbauer, Hakata et al. 2007). To determine whether Vpr directly targets MR using a similar strategy, we co-transfected NL4-3 ΔGPE-GFP or a Vpr-null derivative and an expression vector encoding MR under the control of a cytomegalovirus (CMV) promoter [pCDNA.3.hMR (Liu, Liu et al. 2004)] in HEK293T cells and analyzed MR expression by flow cytometry as shown in Fig S1. We found that Vpr in HEK293T cells had no effect on expression of MR controlled by a heterologous CMV promoter (Figure 2G and S1). Thus, we concluded that Vpr does not degrade MR by the direct mechanism it uses to degrade UNG2 and SMUG1.

In addition to targeting proteins for degradation, Vpr also functions to inhibit transcription of genes such as *IFNA1* (Laguette, Bregnard et al. 2014, Mashiba, Collins et al. 2014). Therefore, we hypothesized that Vpr may reduce MR expression via inhibition of transcription. To examine this, we again utilized infected primary human MDM expressing MR under its native promoter that were transduced with the wild type or Vpr-null reporter virus (Figure 2A). To isolate a uniform population of infected cells necessary for such an analysis, we used fluorescence activated cell sorting to purify transduced (GFP^+^) cells. Indeed, in this system we observed a reduction in MR gene (*MRC1*) expression in transduced (GFP^+^) samples in a Vpr-dependent manner (*p*=0.03, Figure 2H). The magnitude of this effect is consistent with prior reports of HIV-1 inhibiting *MRC1* transcription*−* though this was not previously linked to Vpr (Koziel, Eichbaum et al. 1998, Sukegawa, Miyagi et al. 2018).

### Combined effect of Vpr and Nef dramatically enhances Env levels in primary human MDM

To determine whether the striking downmodulation of MR we observed with expression of both Nef and Vpr affected viral spread in MR^+^ macrophages, we generated additional mutations in HIV-1 89.6 to create a *nef*-null mutant and a *vpr-nef*-null double mutant. As expected, in transfected HEK293T cells these mutations did not alter Env protein levels (Figure 3A) or release of virions as assessed by measuring Gag p24 into the supernatant by ELISAs (Figure 3B). However, in primary human MDM infected with these HIVs, the mutants demonstrated defects in viral spread, with the combination double mutant having the greatest defect (Figure 3C and 3D). The defect in spread was caused in part by diminished virion release, which we previously showed occurred in the absence of Vpr (Mashiba, Collins et al. 2014); MDM infected with the HIV mutants released less Gag p24 even after adjusting for the frequency of infected cells (Figure 3D, right panel), To determine whether the striking downmodulation of MR we observed with expression of both Nef and Vpr affected Env restriction in MR^+^ macrophages, we assessed Env levels in primary human MDM infected with each construct. Because the frequency of infected cells as assessed by intracellular Gag staining (Figure 3C) and Gag pr55 expression as measured by western blot was lower in the mutants than in the wild type infection (Figure 3E), lysate from the wild type sample was serially diluted to facilitate comparisons. Remarkably, we found that the *vpr-nef*-null double mutant, which retains near normal MR levels exhibited the greatest defect in Env expression (Figure 3E, compare lanes with similar Gag as indicated). Mutation of *vpr* and *nef* individually, which greatly increased MR levels compared to wild type HIV infection, also led to striking defects in Env expression. In sum, the effect of Vpr and Nef we observed on MR correlated inversely with Env levels, consistent with MR being the previously reported restriction factor in primary human MDM that restricts Env expression by accelerated lysosomal degradation and which is counteracted by Vpr (Mashiba, Collins et al. 2014). In the case of Vpr, this is likely solely due to downmodulation of MR. In the case of Nef, combined effects on MR and other Env binding proteins including CD4 (Aiken, Konner et al. 1994) and chemokine receptors (Michel, Ganter et al. 2006) may play a role.

**Figure 3:**
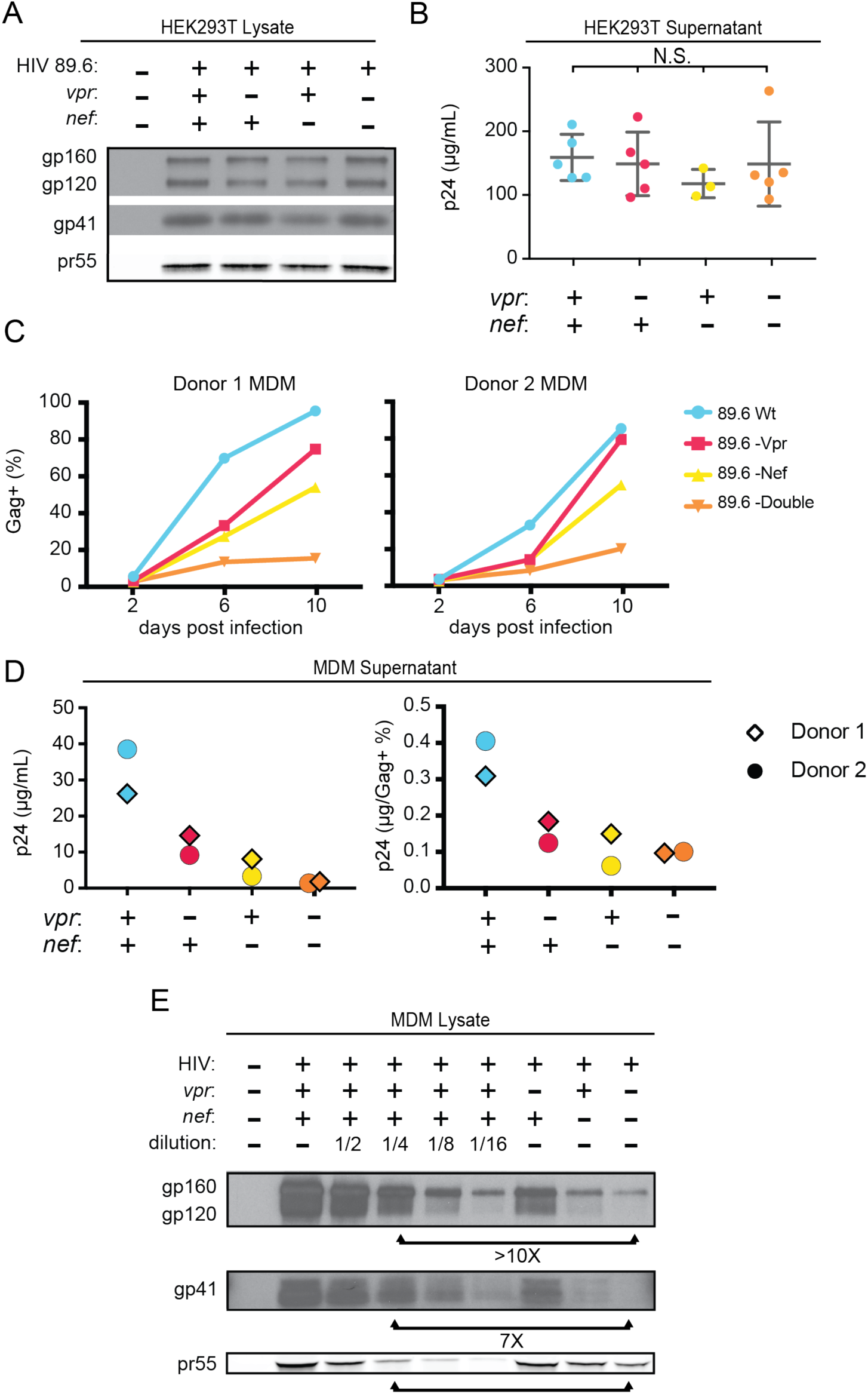
Combined effect of Vpr and Nef dramatically enhances Env levels in primary human MDM. A) Western blot analysis of whole cell lysate from HEK293T transfected with the the indicated HIV construct. B) Summary graph of virion release from HEK293Ts transfected as in A as measured by Gag p24 ELISA. (*n* = 5 independent transfections). The mean +/-standard deviation is shown. Statistical significance was determined by one-way ANOVA. (N.S. – not significant) C) Frequency of infected primary human MDM infected with the indicated HIV and analyzed over time by flow cytometric analysis of intracellular Gag. (For parts C-E, *n*= 2 independent donors). D) Virion release by primary human MDM infected with the indicated HIV and analyzed by Gag p24 ELISA 10 days post infection. In the right panel, virion release was adjusted for frequency of infected cells as measured in part C. E) Western blot analysis of whole cell lysate from primary human MDM infected with the indicated HIV. Lanes 2 - 6 are a serial dilution series of the wild type sample. The arrows below the Gag pr55 bands indicate the dilution (4 fold) of wild type that has approximately the same amount of Gag pr55 as the *vpr*-*nef*-null double mutant.

### Evidence for a role for mannose-containing glycan in restriction of Env in primary human MDM

Previously reported interactions between HIV Env and MR are believed to occur via mannose-containing glycans on Env. Interestingly, the macrophage tropic strain YU-2, which was isolated from the CNS of an AIDS patient (Li, Kappes et al. 1991), lacks a glycan structure known as the mannose patch. This structure is the target of several broadly neutralizing antibodies including 2G12, to which YU-2 is highly resistant (Trkola, Purtscher et al. 1996). We hypothesized that loss of the mannose patch would decrease interactions with MR and reduce the requirement for Vpr to counteract MDM-specific restrictions to virion release and Env expression. To test this hypothesis we examined the extent to which virion release and Env expression were influenced by Vpr in primary human MDM infected with YU-2 or 89.6 HIVs. Remarkably, we observed no significant difference in Gag p24 release between wild type and *vpr*-null YU-2 infection of MDM (Figure 4A). Moreover, the *vpr*-null mutant of YU2 displayed only a minor defect in Env expression compared to Vpr null versions of 89.6 and NL4-3 (Figure 4B).

**Figure 4:**
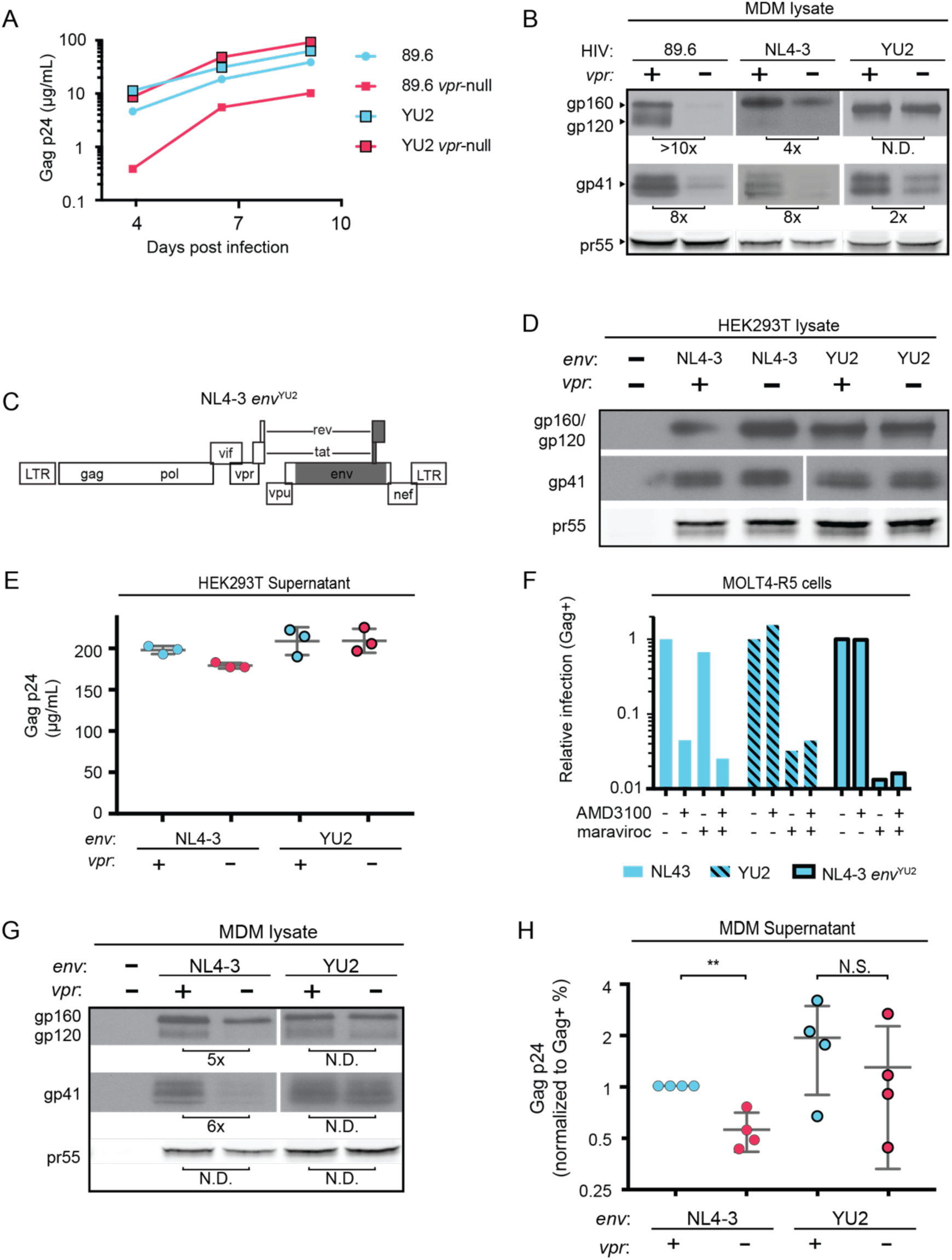
HIV YU2, which lacks a mannose rich patch, does not require Vpr for robust Env protein expression and spread in MDM. A) Virion release over time by primary human MDM infected with the indicated HIV as measured by ELISA (*n*=2 independent donors). B) Western blot analysis of whole cell lysate from MDM infected for 10 days with the indicated HIV. Because NL4-3 infects MDM poorly, NL4-3 was pseudotyped with a YU-2 Env expression plasmid co-transfected in the producer cells as described in Methods. Subsequent spread was blocked in all samples by the addition of entry inhibitors AMD3100 and maraviroc 48 hours post-infection. C) Diagram of the HIV NL4-3 genome. The shaded portion represents the sequence that was replaced with sequence from HIV YU2 to create the NL4-3 *env*^YU-2^ chimera. D) Western blot analysis of HEK293T cells transfected with the indicated HIV constructs. E) Virion release from HEK293T transfected as in E as measured by p24 ELISA. (*n*=3 experimental replicates). F) Relative infection of MOLT4-R5 cells 48 hours after inoculation by the indicated viruses and treated with entry inhibitors as indicated. The frequency of infected cells was measured by intracellular Gag stain and normalized to the untreated condition for each infection. G) Western blot analysis of primary human MDM infected for 10 days with the indicated virus as in B. (*n*=2 independent donors) H) Summary graph showing virion release as measured by p24 ELISA for primary human MDM infected as in G. Virus production was adjusted for infection frequency as determined flow cytometrically using an intracellular Gag stain. The mean +/-standard deviation is shown. (*n*=4 independent donors). N.D. – no difference. Statistical significance was determined using a two-tailed, paired *t*-test. N.S. – not significant, ** p<0.01.

Because there are a number of other genetic differences between YU-2 and the other HIVs, we constructed a chimeric virus, which restricted the differences to the *env* open reading frame. As shown in Figure 4C, a fragment of the YU-2 genome containing most of *env* but none of *vpr* (Figure 4C, shaded portion) was cloned into NL4-3 and NL4-3 *vpr*-null. As expected, these genetic alterations did not affect Env protein levels or virion release in transfected HEK293T cells (Figure 4D and E). To confirm that the chimeric Env was still functional, we examined infectivity in T cells prior to performing our analyses in primary human MDM. Conveniently, sequence variation within the gp120 region allows YU-2 Env to only utilize the co-receptor CCR5 for entry, whereas NL4-3 can only utilize CXCR4. Thus, we expected the NL4 3*env*^YU2^ chimera would switch from being CXCR4- to CCR5-tropic. To test this, we utilized a T cell line expressing both chemokine receptors (MOLT4-R5) and selectively blocked entry via CXCR4 and CCR5 entry inhibitors [AMD3100 and maraviroc, respectively (Figure 4F)]. As expected, entry of MOLT4-R5 cells by NL4-3 was blocked by AMD3100 but not maraviroc, indicating CXCR4-tropism. The chimeric NL4-3 *env*^YU2^ and wild type YU-2 demonstrated the reverse pattern, indicating CCR5-tropism. These results demonstrated that we had made the expected changes in the chimeric Env without disrupting its capacity to infect cells.

To determine whether swapping a limited portion of YU-2 containing Env into NL4-3 alleviated the requirement for Vpr, we examined Env expression and virion release in primary human MDM infected with these viruses. Because the parental NL4-3 virus required pseudotyping with a macrophage-tropic Env for entry and was unable to spread in MDM, all infections were treated with entry inhibitors AMD3100 and maraviroc 48 hours after inoculation to block subsequent rounds of infection. Remarkably, we observed that wild type NL4-3 Env but not chimeric NL4-3 *env*^YU2^ required Vpr for maximal expression (Figure 4G). Moreover, MDM infected with the chimeric HIV had a reduced requirement for Vpr for maximal virion release (Figure 4H). This experiment provides strong evidence that the requirement for Vpr can be alleviated by genetic changes within the *env* open reading frame. These results are consistent with a model in which YU-2 *env* confers resistance to the effects of MR due to the absence of the mannose rich structure on the YU-2 Env glycoprotein.

### Deletion of N-linked glycosylation sites in Env reduces Env restriction in HIV-1 infected human primary MDM and diminishes the need for Vpr and Nef

To more directly assess the role of mannose in restricting expression of Env in HIV-1 infected primary human MDM, we engineered a version of 89.6 Env in which two N-linked glycosylation sites, N230 and N339 (HIV HxB2 numbering) were substituted for amino acids found at analogous positions in YU-2 Env (Figure 5A). The glycosylation sites N2330 and N339 were selected because they contain high-mannose glycan structures (Leonard, Spellman et al. 1990) that are absent in YU-2 Env. Loss of N230 limits neutralization by glycan specific antibodies (Huang, Kang et al. 2014). Loss of N339 decreases the amount of oligomannose (Man_9_GlcNAc_2_) present on gp120 by over 25%, presumably by opening up the mannose patch to processing by *α*-mannosidases (Pritchard, Spencer et al. 2015). These substitions (N230D and N339E) in 89.6 did not alter virion production (Figure 5B) or Env protein expression (Figure 5C) in transfected HEK293T cells.

**Figure 5:**
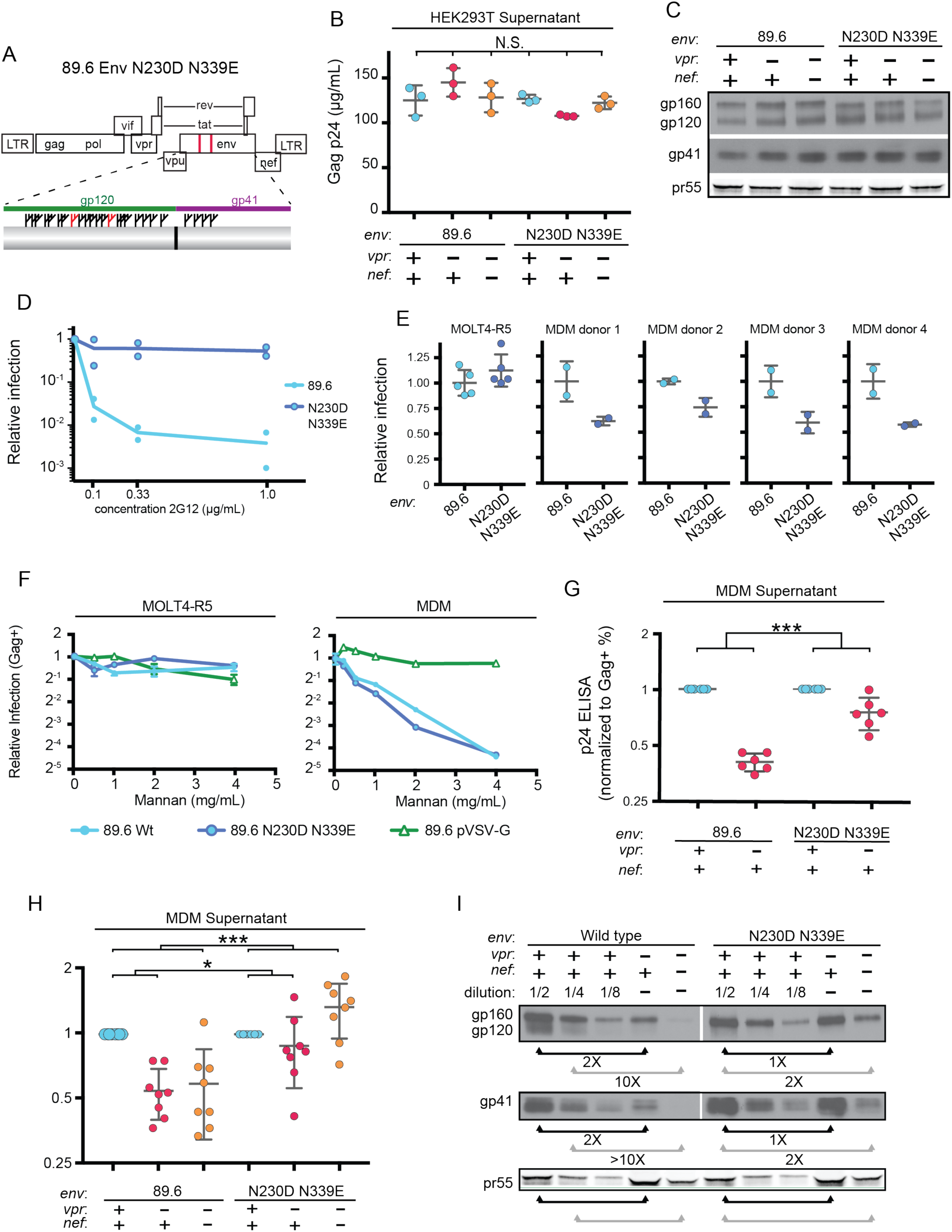
Deletion of N-linked glycosylation sites in *env* reduces the requirement for Vpr and Nef for virion release and Env expression in HIV-1 infected primary human MDM. A) Upper panel, diagram of HIV genome encoding the mutations N230D and N339E (indicated in red) to prevent N-linked glycosylation at those sites. Lower panel, diagram of HIV 89.6 N230D N339E mutant Env protein. Branched symbols represent N-linked glycans. B) Summary graph showing virion release from HEK293Ts transfected with the indicated HIV construct as measured by p24 ELISA. (*n*=3 experimental replicates). Statistical significance was determined by one-way ANOVA. C) Western blot analysis of HEK293T transfected as in B. D) Summary graph showing relative infection frequency of MOLT4-R5 T cells by the indicated HIV following treatment as indicated with the neutralizing antibody 2G12. The percentage of infected cells was measured by intracellular Gag stain and normalized to the untreated condition for each virus. (n= 2 independent experiments, both are plotted) E) Summary graphs of relative infection of the indicated cell type by mutant or parental wild type HIV. The frequency of infected cells was measured flow cytometrically by intracellular Gag stain and normalized to the wild-type virus. (*n*=5 experimental replicates for MOLT4-R5; *n*=2 experimental replicates for MDM from 4 independent donors). F) Summary graph depicting relative infection of the indicated cell type by each virus plus or minus increasing concentrations of mannan as indicated. The frequency of infected cells was measured by intracellular Gag stain and normalized to the uninhibited (0 mg/mL mannan) condition for each virus. 89.6 pVSV-G indicates 89.6 Δenv pseudotyped with VSV-G protein. (*n*=2 independent donors for 89.6 wild type and 89.6 Δenv pVSV-G; *n*=1 donor for 89.6 *env* N230D N339E) G) Summary graph of virion release from primary human MDM following 10 days of infection by the indicated HIV as measured by p24 ELISA. Virion release was normalized to the infection frequency assessed flow cytometrically by intracellular Gag stain. The result for each *vpr*-null mutant was normalized to the *vpr*-competent virus encoding the same *env*. (*n*=6 independent donors) H) Summary graph of virion release from primary human MDM following 10 days of infection by the indicated HIV as measured by p24 ELISA. Virion release was normalized to the infection frequency assessed flow cytometrically by intracellular Gag stain. For this single round infection assay, all viruses were pseudotyped with YU2 Env and viral spread was blocked 48 hours later by addition of AMD3100 and maraviroc. (*n*=8 independent donors) The result for each *vpr*-null or *vpr-nef*-null mutant was normalized to the *vpr*- and nef-competent virus encoding the same *env*. I) Western blot analysis of MDM infected as in E. The lysates from the *vpr*-competent and *nef*-competent infections were diluted to facilitate comparisons to *vpr*- and *nef*-null mutants. (*n*=2 independent donors) For summary graphs, the means +/-standard deviation is shown. In panels G and H statistical significance was determined by a two-tailed, paired *t*-test * p=0.01, ** p<0.01, *** p<0.001.

To confirm that mutation of N230 and N339 disrupted the mannose patch on Env, we assayed the ability of 2G12, which recognizes epitopes in the mannose patch (Sanders, Venturi et al. 2002, Scanlan, Pantophlet et al. 2002), to neutralize wild type and mutant Env. As shown in Figure 5D, wild type but not mannose deficient N230D N339E was neutralized by 2G12. In addition, we found that these substitutions did not disrupt infection of a T cell line that does not express MR (Figure 5E). However, somewhat unexpectedly, we found that HIV containing the N230D N339E Env substitutions were approximately 40% less infectious to primary human macrophages expressing MR than the wild type parental virus (Figure 5E). This macrophage-specific difference in infectivity suggested that mannose on Env facilitates initial infection through interactions with MR, which is highly expressed on differentiated macrophages. To examine this possibility further, we asked whether mannan, which competitively inhibits MR interactions with mannose containing glycans (Shibata, Metzger et al. 1997), was inhibitory to HIV infection. As a negative control, we tested 89.6 Δenv pseudotyped with vesicular stomatitls virus G-protein Env (VSV-G) which has only two N-linked glycosylation sites, both of which contain complex-type rather than high-mannose glycans (Reading, Penhoet et al. 1978) and therefore should not bind MR or be inhibited by mannan. As expected, we found that infection of a T cell line lacking MR was not sensitive to mannan (Figure 5F, left panel). However, infection of MDM by wild type HIV-1 was inhibited up to 16-fold by mannan. This was specific to HIV Env because mannan did not inhibit infection by HIV lacking *env* and pseudotyped with the heterologuos VSV-G Env (Figure 5F). Interestingly, mannan also inhibited baseline macrophage infection by mannose-deficient Env (89.6 Env N230D N339E), indicating that N230D N339E substitutions did not completely abrogate glycans on Env that are beneifical to initial infection. In sum, our results demonstrate that interactions with mannose binding receptors are advantageous for initial HIV infection of macrophages and that the glycans remaining on Env N230D N339E retain some ability to bind glycan receptors on macrophages that facilitate infection.

While interactions between high-mannose residues on Env and MR are advantageous for viral entry, we hypothesized that they interfered with intracellular Env trafficking and were deleterious to egress of Env-containing virions in the absence of Vpr and/or Nef. To test this, we examined virion release and Env expression by HIVs encoding the mannose-deficient Env N230D N339E plus or minus Vpr expression. We found that mannose-deficient Env had a reduced requirement for Vpr for maximal virus relase compared with the parental wild type virus in a spreading infection system (Figure 5G, *p*<0.001). In addition, the mannose-deficient Env had a reduced requirement for both Nef and Vpr in virion release assays using primary human MDM infected for a single round of infection (Figure 5H, p<0.001). Single round infection assays were used to assess the *vpr-nef* double mutant because depletion of mannose on Env did not rescue spread. This is likely due to pleiotropic effects of Nef that disrupt interference by the HIV receptors, CD4, CXCR4 and CCR5 (Lama, Mangasarian et al. 1999, Michel, Allespach et al. 2005, Venzke, Michel et al. 2006) combined with the reduced infectivity of the mannose deficient Env.

Finally, we asked whether the mannose-deficient Env had increased stability in primary human MDM lacking Vpr and/or Nef by western blot analysis. Remarkably, we found that the mannose-deficient Env rescued Env expression in the absence of Vpr (Figure 5I, right side, black bars), and reduced the defect observed in the *vpr*-*nef*-null double mutant (Figure 5I, right side, gray bars) once differences in infection frequency were accounted for by matching pr55 expression in the dilution series. These data provide strong support for the model that MR restricts Env expression via direct interaction with high-mannose residues on Env and that this restriction is counteracted by Vpr and Nef.

### Silencing MR alleviates restriction of Env in primary human MDM lacking Vpr

To directly test the hypothesis that MR is the restriction factor in MDM that is counteracted by Vpr, we examined the effect of MR silencing on Env expression in HIV-infected MDM lacking Vpr. Remarkably, we observed that silencing MR markedly reduced Env restriction - once differences in infection frequency as assessed by Gag pr55 expression were accounted for (Figure 6A). These results support the conclusion that the Env restriction observed in *vpr*-null 89.6 is dependent on expression of MR.

**Figure 6:**
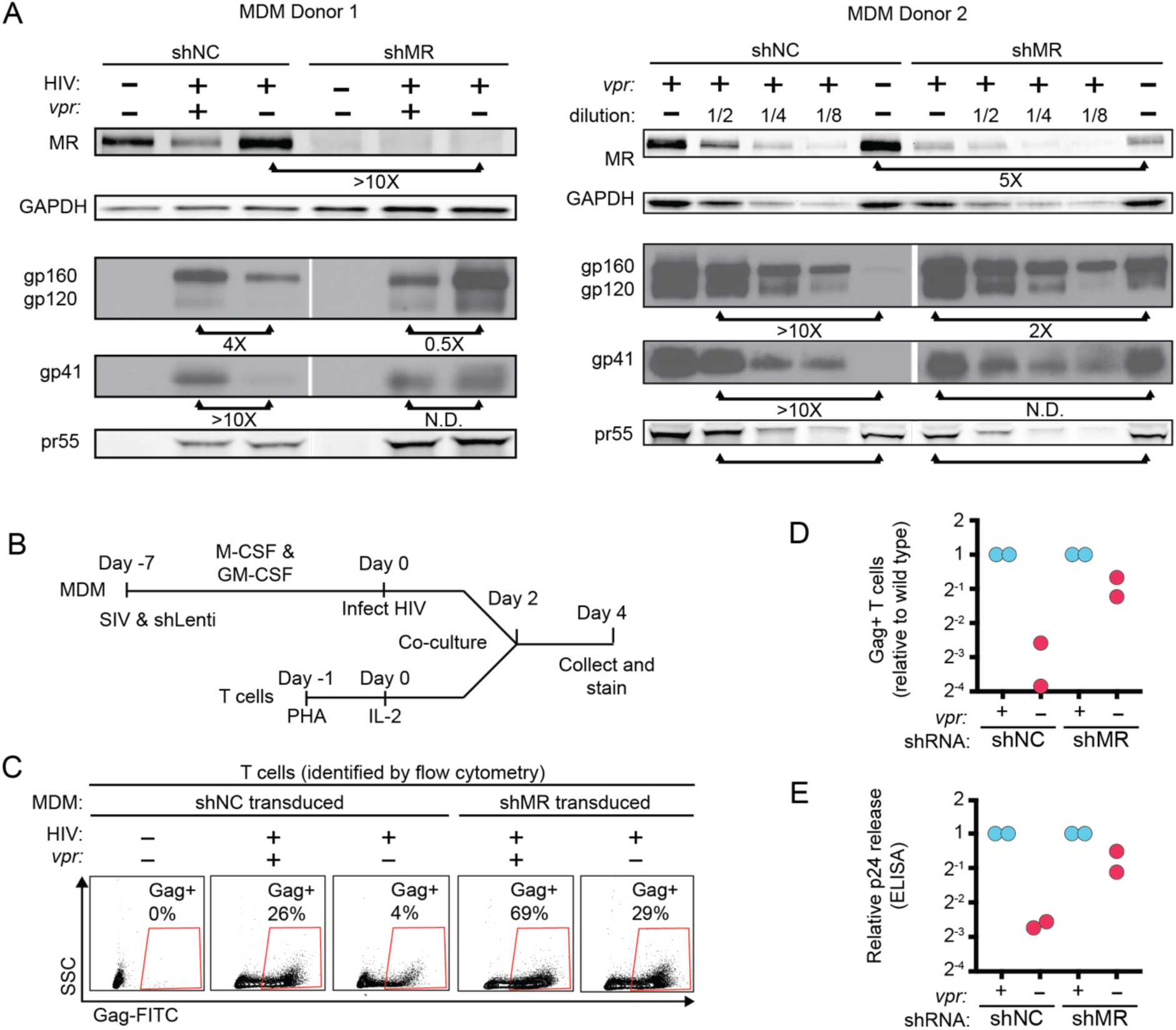
Knockdown of MR enhances Env expression and spread to T cells in *vpr*-null infection of MDM. A) Western blot analysis of MDM from two independent donors treated with the indicated silencing vector and infected with the indicated HIV for 10 days. The miRNA sequences encoded by the negative control vector (shNC) and the MR silencing vector (shMR) are described in Methods. B) Schematic diagram of experimental protocol used for silencing experiments. C) Representative flow cytometric plots showing frequency of infected (Gag^+^) primary T cells following two days of co-culture with autologous, HIV 89.6 infected primary MDM. T cells were identified in co-culture by gating on CD3^+^ CD14^-^ cells as shown in Fig S2. D) Summary graph displaying relative infection of T cells as measured in C and normalized to wild type. (*n*=2 independent donors) E) Virus release by co-cultured MDM and T cells as measured by p24 ELISA and normalized to wild type. (*n*= 2 independent donors).

Previous work in our lab demonstrated that restriction of Env in primary human MDM disrupted formation of virological synapses and cell-to-cell spread of HIV from infected MDM to T cells (Collins, Lubow et al. 2015). Expression of Vpr alleviated these effects, dramatically increasing viral transmission – especially under conditions of low initial inoculum of free virus. To determine whether MR was responsible for these defects in MDM lacking Vpr, we measured Vpr-dependent HIV-1 spread from primary human MDM silenced for MR to activated primary T cells freshly isolated from the same donor. In this assay system, co-cultured cells were stained for CD3 to distinguish T cells and CD14 to distinguish MDM as shown in Figure S2. Indeed, we found that silencing MR dramatically reduced the requirement for Vpr to support spread from MDM to T cells (Figure 6D). In addition, MR silencing reduced the need for Vpr in virus release assays in the co-culture system (Figure 6E). These data support the conclusion that MR is the previously identified restriction factor in macrophages that reduces spread from Vpr-null HIV-infected macrophages to T lymphocytes.

### Infection of macrophages dramatically facilitates intial infection by transmitter/found (T/F) HIVs involved in intial HIV transmission

Because macrophages are present in genital mucosa, are permissive to HIV infection (Shen, Richter et al. 2009), and recruit HIV-permissive CD4^+^ T cells as part of their immunologic function (Liao, Rabin et al. 1999) we wondered whether cell-to-cell spread from MDM to T cells may potentially facilitate transmission. To examine this possibility, we tested two transmitted/founder (T/F) HIV molecular clones (REJO and CH077), which can both cause detectable infection in T cells following spinnoculation (Figure 7A, upper panel) or long term culture (Ochsenbauer, Edmonds et al. 2012), but which differ in their capacity to infect macrophages (Ochsenbauer, Edmonds et al. 2012). We observed that when MDM were inoculated briefly (6 hours) and cultured for two days, REJO produced detectable infection but CH077 did not (Figure 7A, lower panel). Using the experimental protocol diagrammed in Figure 7B, we observed that when T cells were cultured for two days with cell free virus without spinnoculation, both T/F viruses failed to infect a significant fraction of T cells (Figure 7C, upper panel). In contrast, when T cells were co-cultured for two days with MDM that had previously been infected as in Figure 6F, the T/F virus REJO was transmitted to T cells but CH077 was not (Figure 7C, lower panel) Thus, under conditions of limited exposure to cell free virus, macrophage infection can dramatically enhance spread and speed infection to T cells. The role for Vpr in this process likely contributes to its evolutionary conservation in nearly all simian and human lentivirus genomes (Mashiba and Collins 2013).

**Figure 7:**
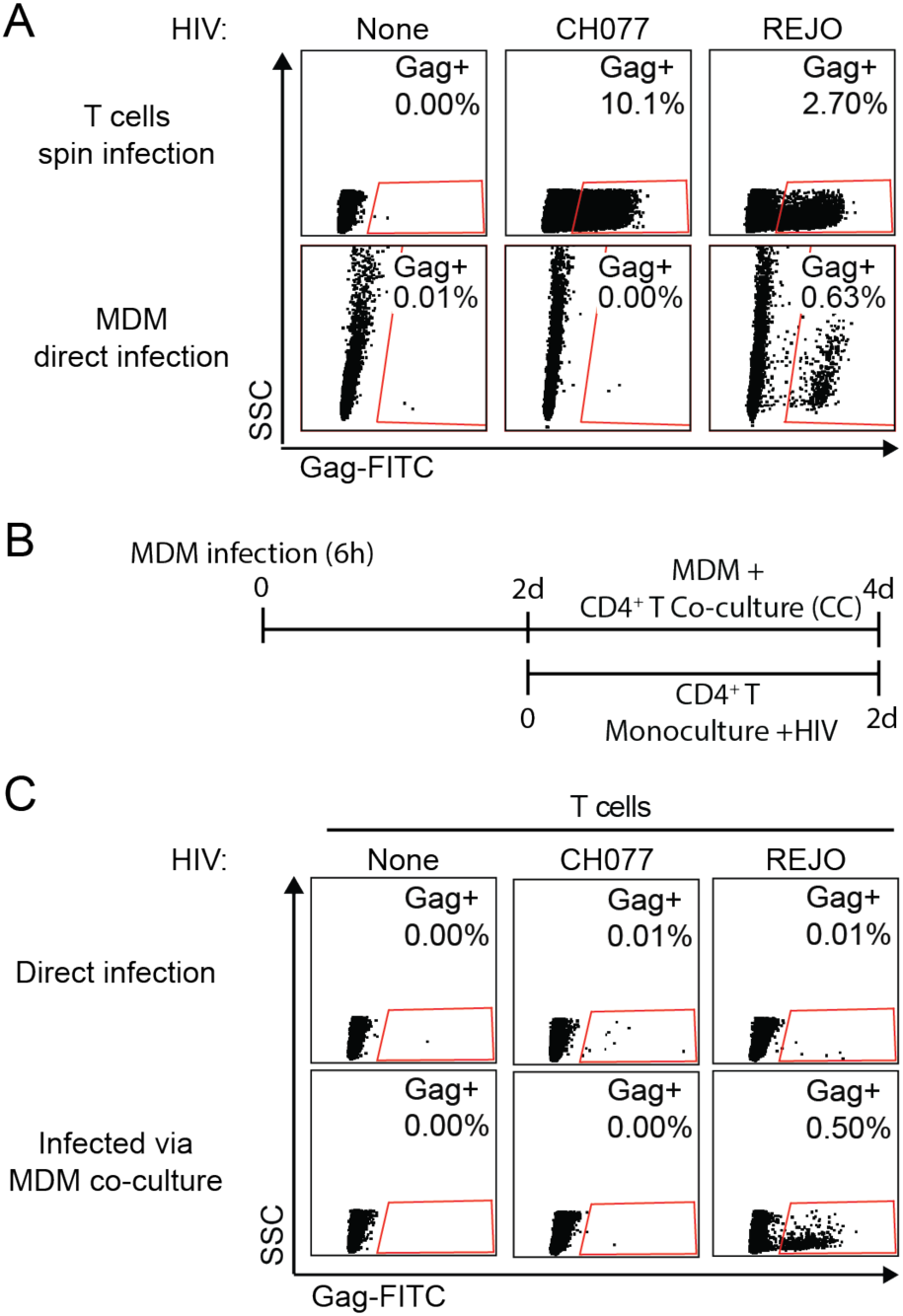
MDM enhance infection of T cells by transmitted/founder (T/F) viruses. A) Flow cytometry plots of primary human T cells 2 days after 2 hour spinnoculation or MDM 4 days after six hour incubation with 50µg of the T/F virus indicated. B) Schematic diagram of protocol for experiments shown in part C. C) Flow cytometry plots to measure the percentage of Gag^+^ T cells 2 days after infection by the method indicated. For direct infection, T cells were continuously cultured with with 50µg of virus over the two day incubation period. For infection via MDM co-culture, T cells were co-cultured with MDM transiently exposed to 50µg of virus as in part A. For the co-culture assays, T cells were identified in co-culture by gating on CD3^+^ CD14^-^ cells. *n*=1 donor.

## Discussion

Our findings demonstrate that HIV encodes two accessory proteins, Vpr and Nef, that dramatically reduce expression of MR on infected macrophages and provide an explanation for the evolutionary conservation of Vpr in HIV-1. Our results clearly define MR as a factor capable of restricting the expression of Env and interfering with virion release. While the effect of Nef on MR was previously reported, the effect of Vpr on MR was entirely unexpected. The evidence that Vpr counteracts MR to stabilize Env is strong. We demonstrate that Vpr expression correlates with reduced MR expression. In addition, we show that loss of Vpr can be rescued by changes within the Env locus that reduce detrimental interactions with MR, including point mutations that selectively delete mannose residues. Finally, silencing MR expression decreased the requirement for Vpr to stabilize Env and enhanced spread to T cells and virion release.

Other investigators have reported that MR inhibits virion egress in HEK293T cells (Sukegawa, Miyagi et al. 2018). However, this study differed from results we report here in primary human macrophages because the prior study observed effects on virions that were Env-independent. In addition, they did not examine effects of Vpr on MR in their system (Sukegawa, Miyagi et al. 2018). In the primary macrophage system, Vpr-senstiive virion restriction depends entirely on an intact *env* open reading frame (Mashiba, Collins et al. 2014) and genetic changes in the *env* open reading frame – especially those that alter N-linked glycosylation sites -critically affect the requirement for Vpr. The effect of MR on Env and Env-containing virion release helps explain the significance behind prior in vivo observations demonstrating that HIV infection reduces expression and activity of MR in infected humans (Koziel, Kruskal et al. 1993, Koziel, Eichbaum et al. 1998) and that SIV does the same in monkeys (Holder, McGary et al. 2014). By demonstrating how the effect of Vpr on MR promotes macrophage to T cell spread, we also provide an explanation for how Vpr increases infection of human lymphoid tissue *ex vivo* (Eckstein, Sherman et al. 2001, Rucker, Grivel et al. 2004), which contain macrophages and T cells in a highly physiological, three-dimensional environment.

As Nef had already been shown to reduce MR surface expression (Vigerust, Egan et al. 2005), the observation that HIV encodes a second protein, Vpr, to reduce MR expression was unanticipated, but not unprecedented; other host proteins are known to be affected by more than one lentiviral accessory protein. The HIV receptor, CD4, is simultaneously targeted by Vpu, Nef and Env in HIV-1 (Chen, Gandhi et al. 1996) and tetherin is alternately targeted by Vpu, Nef, or Env in different strains of primate lentiviruses (Harris, Hultquist et al. 2012). Nef has also been shown to downmodulate the viral co-receptors CXCR4 (Venzke, Michel et al. 2006) and CCR5 (Michel, Allespach et al. 2005), which may also interfere with Env expression in infected cells. Nef’s activity against CXCR4, CCR5, and MR presumably has the same ultimate purpose as its activity against CD4, namely to stabilize Env, enhance virion release and prevent superinfection of the producer cell (Lama, Mangasarian et al. 1999, Ross, Oran et al. 1999). The need for Vpr and Nef to simultaneously target MR may be explained by the high level of MR expression, estimated at 100,000 copies per macrophage (Stahl, Schlesinger et al. 1980). Alone, Vpr and Nef each has a modest effect on MR in primary human macrophages. The magnitude of the effect and the fraction of cells affected increased when both proteins were expressed. The potent combined effect likely derives from synergistic targeting of MR at two different stages of MR synthesis. Nef was shown to alter MR trafficking (Vigerust, Egan et al. 2005) and we show Vpr inhibits MR transcription.

In sharp contrast to the effect we observed in MDM, Vpr did not affect MR protein levels when MR was expressed via a heterologous promoter in the HEK293T cell line, which is derived from human embryonic kidney cells, which are not a natural target of HIV. The cell type selectivity in these experiments is likely due to differences in the promoters driving MR expression, however, we cannot rule out the existence of other macrophage specific pathways required to recreate the effect of Vpr on MR. Further work will be needed to examine these questions and determine other mechanistic details. For example, it will be important to determine whether Vpr utilizes its cellular co-factors to affect MR levels. Vpr is perhaps best known for its interaction with Vpr binding protein [DCAF1, (McCall, Miliani de Marval et al. 2008)], a component of the cellular DCAF1-DDB1-CUL4 E3 ubiquitin ligase complex that plays a role in Vpr-dependent disruption of the cell cycle and cellular DNA repair pathways in dividing cells (Belzile, Duisit et al. 2007, Hrecka, Gierszewska et al. 2007, Le Rouzic, Belaidouni et al. 2007, Wen, Duus et al. 2007, Lahouassa, Blondot et al. 2016, Wu, Zhou et al. 2016, Zhou, DeLucia et al. 2016). This interaction has also been linked to transcriptional inhibition of type I interferons in response to infection in macrophage cultures (Laguette, Bregnard et al. 2014, Mashiba, Collins et al. 2014). Additional research is now needed to determine whether repression of MR and IFN transcription is mediated by the same DCAF-1 dependent pathway.

Interactions between MR and Env are likely mediated by the unusually high density of N linked glycosylation sites that retain high-mannose glycans, which is a known PAMP (Stahl and Ezekowitz 1998, McGreal, Rosas et al. 2006). Here, we show that selective deletion of mannose residues alleviated the requirement for Vpr. Deletion of individual glycosylation sites is known to lead to changes in the processing of neighboring glycans and deletions at certain sites lead to larger than expected losses of oligomannose (Balzarini 2007) presumably because their removal allows greater access and facilitates trimming of surrounding glycans. Selective pressure to maintain mannose residues on Env may be due to the enhanced attachment they mediate. Indeed, we provide strong evidence that Env’s interaction with MR boosts initial infection of MDM. This finding is supported by a prior report that MR enhances HIV-1 binding to macrophages and transmission of the bound virus to co-cultured T cells (Nguyen and Hildreth 2003). Our study adds to these findings by providing evidence that interactions with mannose binding receptors also enhance direct infection of macrophages. Moreover, the capacity of Vpr and Nef to mitigate the effect of detrimental intracellular interactions during viral egress limits the negative impact of retaining high-mannose on Env. In addition, the dense glycan packing, which is privileged from antibody recognition through immune tolerance, is believed to play a role in evasion of the antibody response (Stewart-Jones, Soto et al. 2016).

Here we also confirm our prior observation that Vpr enhances transmission of HIV from infected macrophages to primary T lymphocytes. Macrophages prepared according to our culture conditions are more easily infected by cell free virus in vitro than activated T cells, which require spinoculation for detectable infection following short incubations (48 hours). The observation that T/F viral strains with a greater capacity to infect macrophages have an advantage in our co-culture system suggests a role for Vpr during transmission. The accelerated spread to T cells we observed may be critical to establishing a persistent infection before innate and adaptive immune responses are activated. While in vitro assays of T/F viruses have suggested strong evolutionary selection for T cell infection with limited capacity to infect macrophages, other studies indicate that T/F viruses are generally less infectious to all cell types compared to viruses isolated at other stages (Peters, Gonzalez-Perez et al. 2015), but appear to be selected primarily by their resistance to interferon (Iyer, Bibollet-Ruche et al. 2017). The strong selective pressure to retain Vpr despite its limited effect on T cell-only cultures indicates there is more to learn about the role of Vpr, macrophages and T/F viruses in HIV transmission and pathogenesis. Collectively, these studies suggest that novel therapeutic approaches to inhibit the activity of Vpr and Nef in macrophages would potentially represent a new class of antiretroviral drug that could be an important part of a treatment or prophylactic cocktail.

## Materials and Methods

### Viruses and viral vectors

The following molecular clones were obtained via the AIDS Reagent Program: p89.6 (cat# 3552) from Dr. Ronald G. Collman, pNL4-3 (cat# 114) from Dr. Malcolm Martin, pREJO.c/2864 (cat# 11746) and pCHO77.t/2626 (cat# 11742) from Dr. John Kappes and Dr. Christina Ochsenbauer and pYU2 (cat# 1350) from Dr. Beatrice Hahn and Dr. George Shaw. pCDNA.3.hMR was obtained from Dr. Johnny J. He (Liu, Liu et al. 2004). *Vpr*-null versions of HIV molecular clones were created by cutting the AflII site within *vpr* and filling in with Klenow fragment. A *nef*-null version of 89.6 was created by deleting *nef* from its start codon to the XhoI site. To do this, a PCR amplicon was generated from the XhoI site in *env* to *env*’s stop codon. The 3’ reverse primer added a XhoI site after the stop codon. The 89.6 genome and the amplicon were digested with XhoI and ligated together. (5’ primer CACCATTATCGTTTCAGACCCT and 3’ primer TCTCGAGTTTAAACTTATAGCAAAGCCCTTTCCA)

pSIV3+, pSPAX2, pAPPM-1221 were obtained from Dr. Jeremy Luban (Pertel, Reinhard et al. 2011). pSIV3+ *vpr*-null was generated using a synthesized fragment (ThermoFisher Scientific, Waltham, Massachusetts) of the SIV genome in which the Vpr start codon was converted to a stop codon. pYU2 env was obtained from Dr. Joseph Sodroski (Sullivan, Sun et al. 1995). Creation of pNL4-3 ΔGPE-GFP was described previously (McNamara, Ganesh et al. 2012).

### Primary MDM and T cell isolation and culture

Leukocytes isolated from anonymous donors by apheresis were obtained from the New York Blood Center Component Laboratory. The use of human blood from anonymous, de-identified donors was classified as non-human subject research in accordance with federal regulations and thus not subjected to formal IRB review. Peripheral blood mononuclear cells (PBMCs) were purified by Ficoll density gradient. CD14^+^ monocytes were positively selected using a CD14 sorting kit (StemCell Inc., Vancouver, Canada) following the manufacturer’s instructions. Monocyte-derived macrophages (MDM) were obtained by culturing monocytes in R10 [RPMI-1640 with 10% certified endotoxin-low fetal bovine serum (Invitrogen, ThermoFisher)], penicillin (10 Units/ml), streptomycin (10 μg/ml), L-glutamine (292 μg/ml), carrier-free M-CSF (50 ng/ml, R&D Systems, Minneapolis, Minnesota) and GM-CSF (50 ng/ml, R&D Systems) for seven days. Monocytes were plated at 5×10^5^ cells/well in a 24 well dish, except for those to be transduced with lentivirus and puromycin selected, which were plated at 1×10^6^ cells/well.

CD4^+^ T lymphocytes were isolated from PBMCs by CD8 negative selection (DynaBeads, ThermoFisher), cultured in R10 for several days and activated with 5 μg/ml phytohaemagglutinin (PHA-L, Calbiochem, Millipore Sigma, Burlington, Massachusetts) overnight before addition of 500 IU/ml recombinant human IL-2 (R&D Systems).

### Silencing by shRNA

Short hairpin RNA-mediated silencing was performed as previously described (Pertel, Reinhard et al. 2011, Collins, Lubow et al. 2015). Briefly, we spinoculated freshly isolated primary monocytes with VSV-G-pseudotyped SIV3+ *vpr*-null at 2500 rpm for 2 hours with 4 μg/ml polybrene to allow Vpx-dependent degradation of SAMHD1. Cells were then incubated overnight in R10 with M-CSF (50 ng/ml) and GM-CSF (50 ng/ml) plus VSV-G-pseudotyped lentivirus containing an shRNA cassette targeting luciferase (shNC) or MR (shMR). Following an overnight incubation, the cells were cultured for 3 days in fresh medium before addition of 10 μg/ml puromycin for 3 additional days prior to HIV-1 infection. shRNA target sequences used: *Luciferase*: 5’-TACAAACGCTCTCATCGACAAG-3’, *MRC1:* 5’-ATTGATTATCAGTCAAGTTACT-3’

### Virus production

Virus stocks were obtained by transfecting HEK293T cells (ATCC, Manassas, Virginia) with viral DNA and polyethylenimine (PEI). Cells were plated at 2.5×10^6^ cells per 10cm dish and incubated overnight. The following day 12 µg of total DNA was combined with 48 µg of PEI, mixed by vortexing, and added to each plate of cells. For NL4-3ΔGPE-GFP cells were transfected with 4 µg viral genome, 4 µg pCMV-HIV, and 4 µg VSV-G plasmid. For SIV3+ ΔVpr the cells were transfected with 10.5 µg of viral genome and 1.5 µg VSV-G plasmid. Viral supernatant was collected 48 hours post-transfection and centrifuged at 1500 rpm 5 min to remove cellular debris. SIV3+ ΔVpr was pelleted by centrifugation at 14,000 rpm for 4 hours at 4°C and resuspended at 10x concentration. Virus stocks were aliquoted and stored at -80°C.

### Co-transfections

Co-transfections of HIV and MR were performed in HEK293T cells. Cells were plated at 1.6×10^5^ per well in a 12-well dish. The following day 100 ng of pcDNA.3.hMR, 250 ng of NL4-3 ΔGPE-GFP, and 750 ng pUC19 plasmid was combined with 4.4µg PEI, mixed by vortexing, and added to each well. Cells were lifted using enzyme free cell dissociation buffer (ThermoFisher, cat # 12151014) 48 hours later and analyzed by flow cytometry.

### HIV infections of MDM

Prior to infection, 500µL of medium was removed from each well and this “conditioned” medium was saved to be replaced after the infection. MDM were infected by equal inocula of HIV as measured by Gag p24 mass in 500µL of R10 for 6 hours at 37°C. After 6 hours, infection medium was removed and replaced with a 1:2 mixture of conditioned medium and fresh R10. Where indicated, HIV spread was blocked by AMD3100 (10µg/mL, AIDS Reagent Program cat# 8128) and/or Maraviroc (20µM, AIDS Reagent Program cat# 11580).

### Spin transduction of MDM with NL4-3 ΔGPE-GFP

MDM were centrifuged at 2500rpm for 2 hours at 25°C with equal volume of NL4-3 ΔGPE-GFP or an isogenic mutant in 500uL total medium. Following infection, medium was removed and replaced with a 1:2 mixture of conditioned medium and fresh R10.

### Adenoviral transduction of MDM

Adenovirus was prepared by the University of Michigan Vector Core, and the transduction of MDM was performed as previously described (Leonard, Filzen et al. 2011) at an MOI of 1000 based on HEK293T cell infection estimations and the concentration of particles as assessed by OD_280_.

### Infection of T cells

Activated T cells were infected by three methods as indicated. For direct infection, 5×10^5^ cells were plated per well with 50µg HIV p24 in 500µL R10 +100IU/mL of IL-2 and incubated at 37°C for 48 hours. For spin infection, 5×10^5^ cells were plated per well with 50µg HIV and 4ng/mL polybrene in 500µL R10 +100IU/mL of IL-2 and centrifuged at 2500rpm for 2 hours at 25°C. After both types of infection, medium was removed and replaced with 1mL R10 +100IU/mL of IL-2. For co-culture with autologous, infected MDM medium was removed from MDM wells and 5×10^5^ T cells were added in 1mL R10 +100IU/mL of IL-2. All T cell infections were collected 48 hours post infection.

### Flow cytometry

Intracellular staining of cells using antibodies directed against HIV Gag p24 and MR was performed by permeabilizing PFA-fixed cells with 0.1% Triton-X in PBS for 5 min, followed by incubation with PE-conjugated antibody for 20 minutes at room temperature. Surface staining for CD4, CD3 and CD14 was performed before fixation as described previously (Collins, Lubow et al. 2015). Flow cytometric data was acquired using a FACSCanto instrument with FACSDiva collection software (BD, Franklin Lakes, New Jersey) or a FACScan (Cytek, BD) with FlowJo software (TreeStar, Ashland, Oregon) and analyzed using FlowJo software. Live NL4-3 ΔGPE-GFP transduced cells were sorted using a FACSAria III (BD) and gating on GFP^+^ cells.

### Quantitative RT-PCR

MDM sorted as described above in “Flow cytometry” were collected into tubes containing RLT buffer (Qiagen, Hilden, Germany) and RNA was isolated using RNeasy Kit (Qiagen) with on-column DNase I digestion. RNA was reverse transcribed using qScript cDNA SuperMix (Cat #95048, Quantabio, Beverly, Massachusetts). Quantitative PCR was performed using TaqMan Gene Expression MasterMix (ThermoFisher, cat# 4369016) on an Applied Biosystems 7300 Real-Time PCR System using TaqMan Gene Expression primers with FAM-MGB probe. The primers for *ACTB* are assay Hs99999903 and for *MRC1* are assay Hs00267207 (ThermoFisher). Reactions were quantified using ABI Sequence Detection software compared to serial dilutions of cDNA from mock-treated cells. Measured values for *MRC1* were normalized to measured values of *ACTB*.

### Immunoblot

MDM cultures were lysed in Blue Loading Buffer (Cell Signaling Technologies, Danvers, Massachusetts), sonicated with a Misonix sonicator (Qsonica, LLC., Newtown, Connecticut) and clarified by centrifugation at 8000 RPM for 3 minutes. Lysates were analyzed by SDS-PAGE immunoblot. The proteins MR, GAPDH and pr55 were visualized using AlexFluor-647 conjugated secondary antibodies on a Typhoon FLA 9500 scanner (GE, Boston, Massachusetts) and quantified using ImageQL (GE). The proteins gp160, gp120, gp41, Nef, Vpr and GFP were visualized using HRP-conjugated secondary antibodies on film. Immunoblot films were scanned and the mean intensity of each band, minus the background, was calculated using the histogram function of Photoshop CC (Adobe, San Jose, California).

### Virion Quantitation

Supernatant containing viral particles was lysed in Triton X lysis buffer (0.05% Tween 20, 0.5% Triton X-100, 0.5% casein in PBS). CAp24 antibody (clone 183-H12-5C) was bound to Nunc MaxiSorp plates (ThermoFisher) at 4°C overnight. Lysed samples were captured for 2 hr and then incubated with biotinylated antibody to CAp24 (clone 31-90-25) for 1 hr. Clone 31-90-25 was biotinylated with the EZ-Link Micro Sulfo-NHS-Biotinylation Kit (Pierce, ThermoFisher). Samples were detected using streptavidin-HRP (Fitzgerald, Acton, Massachusetts) and 3,3′,5,5′-tetramethylbenzidine substrate. CAp24 concentrations were measured by comparison to recombinant CAp24 standards (ViroGen, Watertown, Massachusetts).

### Antibodies

Antibodies to CAp24 (clone KC57-PE, cat# 6604667, Beckman Coulter, Brea, California), CD3 (clone OKT3-Pacific Blue, cat# 317313, BioLegend, San Diego, California), CD14 (clone HCD14-APC, cat# 325608, BioLegend), CD4 (clone OKT4, cat#17-0048-42, Invitrogen, ThermoScientific), and MR (clone 19.2-PE, cat# 555954, BD) were used for flow cytometry. Antibodies to the following proteins were used for immunoblot analysis: MR (cat# ab64693, Abcam, Cambridge, Massachusetts), GAPDH (clone 3C2, cat# H00002597-M01, Abnova, Taipei, Taiwan), Gag pr55 (HIV-Ig AIDS Reagent Program cat# 3957), Env gp160/120 (AIDS Reagent Program Cat# 288 from Dr. Michael Phelan) Env gp41 (AIDS Reagent Program cat# 11557 from Dr. Michael Zwick), Vpr (AIDS Reagent Program cat# 3951 from Dr. Jeffrey Kopp). Neutralizing antibody 2G12 (AIDS Reagent Program cat# 1476 from Dr. Hermann Katinger) was used at a 1 μg/ml for at the time of infection.

## Acknowledgments

This research was supported by the NIH (5T32GM008353-27 to J.L., R01AI046998 to K.L.C., R56AI130004 to K.L.C., R21AI32379 to K.L.C., T32GM007863 to M.M., T32AI007413 to M.M. and T32AI007528 to D.R.C.). We are grateful to the University of Michigan Vector Core and the NIH AIDS Reagent Program for reagents.

---

**Supplement Figure 1:**
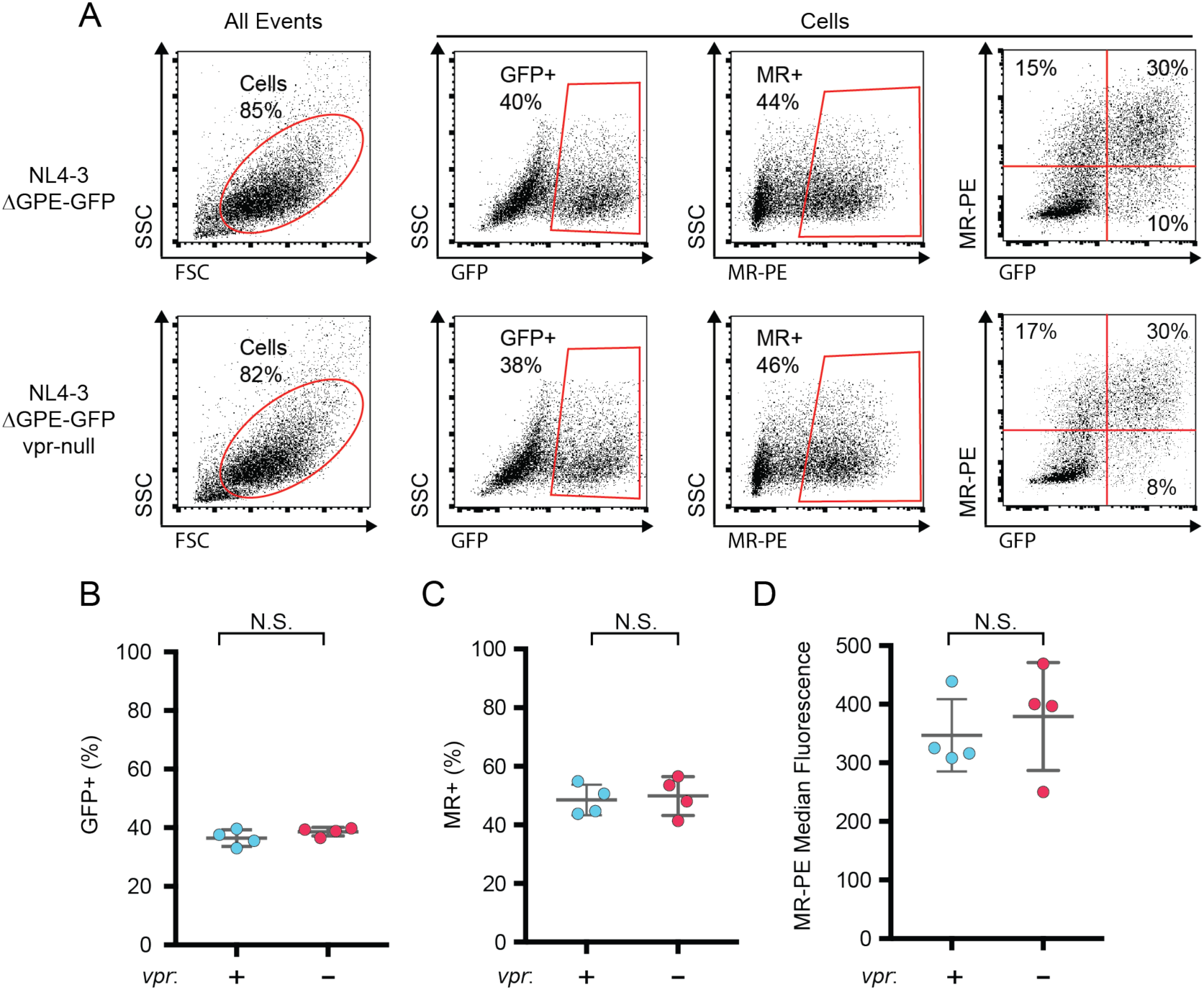
Vpr does not affect MR expression in co-transfected HEK293T cells. A) Representative flow cytometric analysis of HEK293T cells co-transfected with the indicated GFP-expressing HIV genome (NL4-3 ΔGPE-GFP plus or minus an intact *vpr* open reading frame) and an expression vector containing MR (pcDNA3.hMR). The transfections were performed four times and a representative pair of *vpr*-competent and *vpr*-null transfections was chosen. B) Summary graph showing the percentage of cells that are transfected (GFP+) following co-transfection as in A. C) Summary graph showing the percentage of cells that remain MR+ following co-transfection as in B. D) Summary graph showing the the median MR-PE fluorescence of all cells following co-transfection as in A. For all graphs, the mean plus or minus standard deviation is shown. (*n*=4 experimental replicates). Statistical significance was determined by a two-tailed, paired *t*-test. N.S. – not significant: p=0.30, p=0.66, and p=0.57 respectively.

**Supplemental Figure 2:**
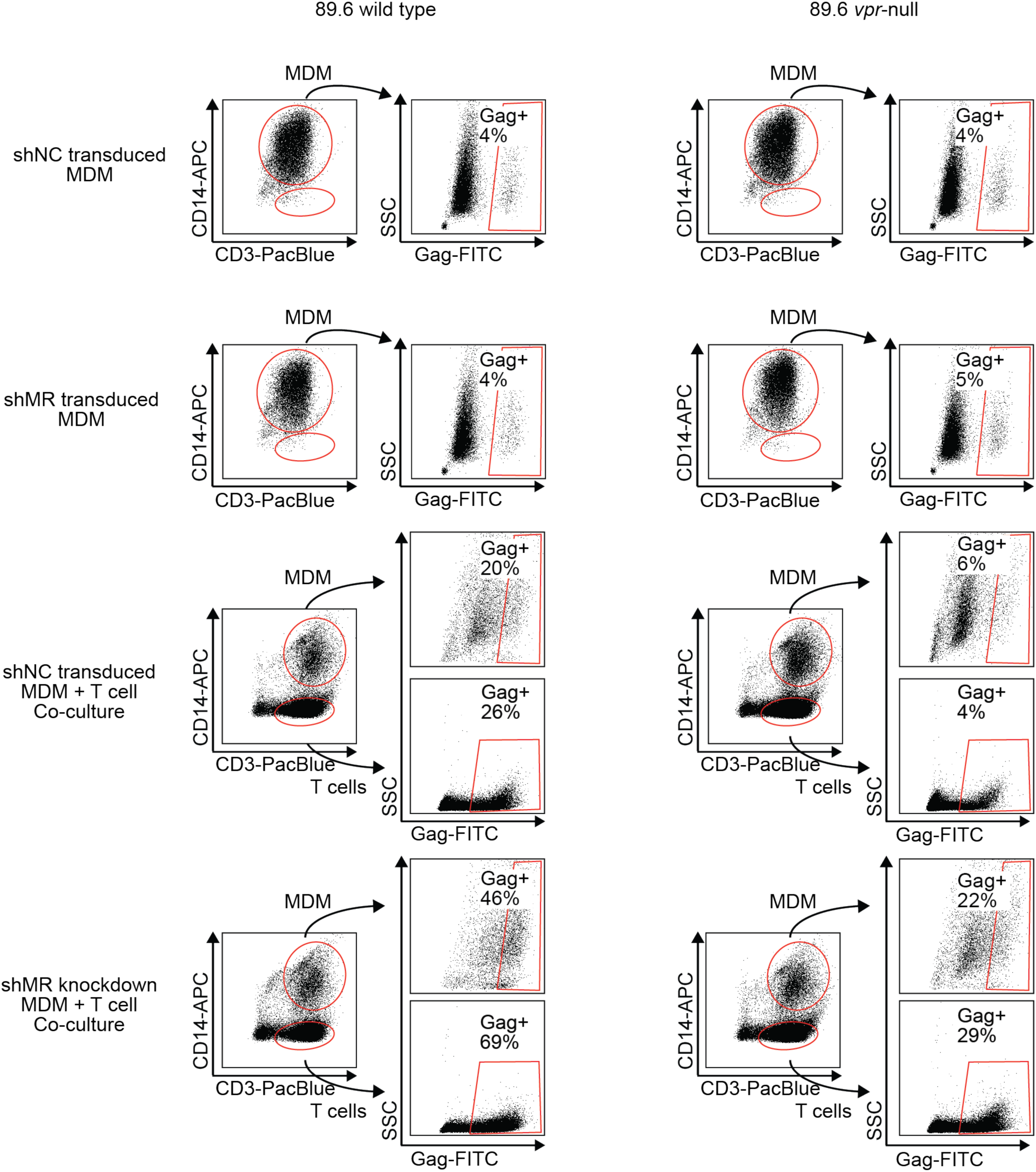
T cells and MDM can be identified by flow cytometry and the percentage of each that are Gag+ can be measured separately. Flow cytometric dot plots illustrating segregation of CD14+ MDM from CD3+ T cells in co-cultures and subsequent assessment of HIV-1 infection by intracellular Gag p24 stain after treatment of the indicated cultures treated as shown in Fig 6B. *n*=2 independent donors. Plots from one donor are shown here and Gag+ percentages from both are shown in Figure 6.

